## Supplemental Material for "Time-resolved oxidative signal convergence across the algae–embryophyte divide"

**SUPPLEMENTARY MATERIAL**

**HPLC-UV-Vis-DAD**

**Method optimization**

**Extraction of and separation of pigments**

The extraction protocol was originally inspired by Aronsson et al 2008. First (as in Aronsson et al 2008) extraction was tested with fresh weight but resulted in chromatograms with asymmetrical signals/insufficient chromatographic separation. Lyophilized material solved this problem. We noticed that protection from light, heat, and oxygen during extraction was essential for reproducible results so we took several measures to do so:
1. All extractions were performed in a dark room (only minimal indirect light) at 4 °C.
2. 0.1 w% BHT were added to the extraction solvents.
3. Extractions were performed swiftly and samples were analyzed directly.
4. Any homogenization of plant material was performed in liquid nitrogen under light shielded conditions.

We started method optimization of separation with an RP-$C_{18}$ column but could not separate lutein and zeaxanthin properly. So, we switched to an RP-$C_{30}$ column where we finally managed after some gradient optimization to separate all carotenoid isomers. The gradient described in the method section of the main publication was based on Gupta et al 2015 but needed to be fine-tuned to reproducibly separate all pigments in diverse lineages and species.

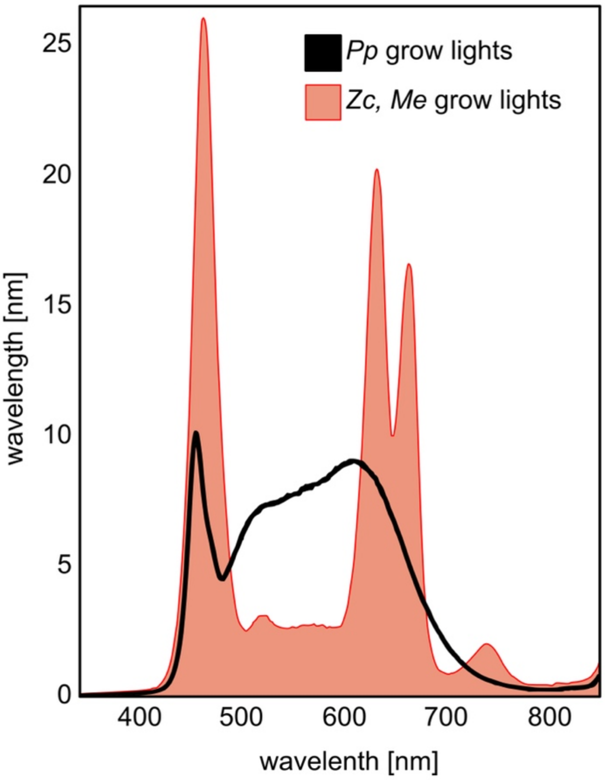


**Supplemental Figure 1:** Spectrum of growth lights used for the cultivation of *Physcomitrium patens* (*Pp*), *Zygnema circumcarinatum* (*Zc*), and *Mesotaenium endlicherianum* (*Me*).

**Representative HPLC chromatogram of *Mesotaenium endlicherianium* and respective absorption spectra**

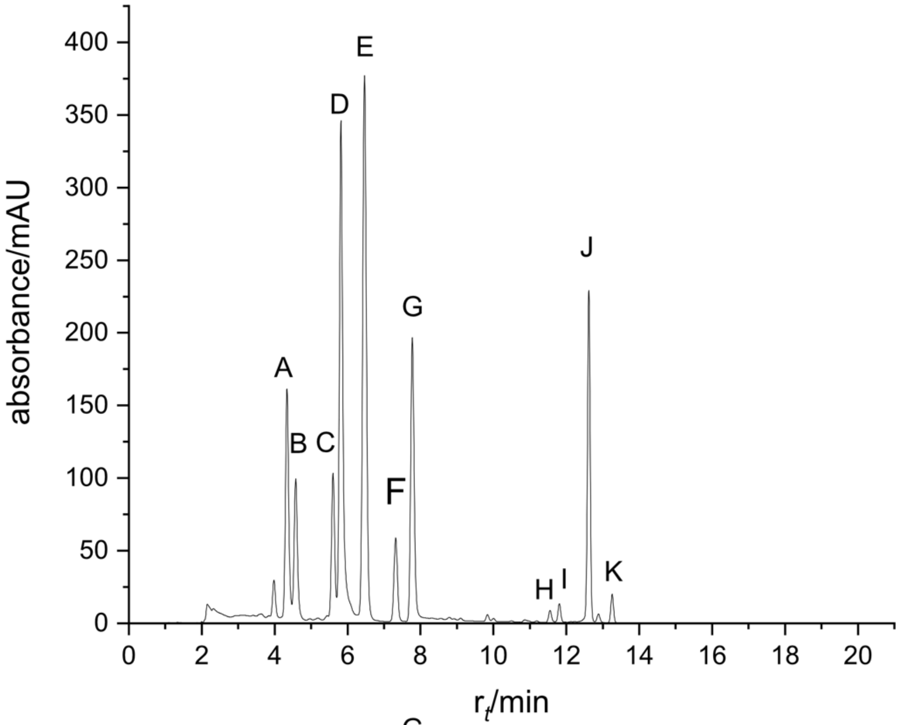


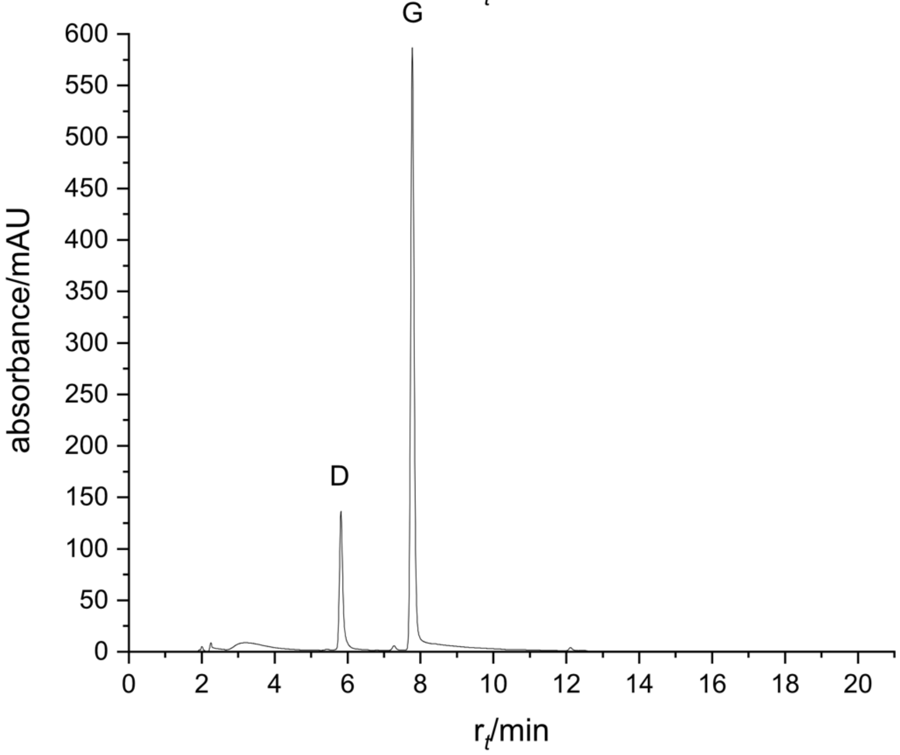


**Supplemental Figure 2:** HPLC chromatograms *Mesotaenium endlicherianum* control replicate 1 recorded at 451 nm (top) and 660 nm (bottom). Violaxanthin (A), 9-*cis*-neoxanthin (B), antheraxanthin (C), chlorophyll b (D), lutein (E), zeaxanthin (F), chlorophyll a (G), 15-*cis*-β-carotene (H), α-carotene (I), β-carotene (J), 9-*cis*-β-carotene (K).

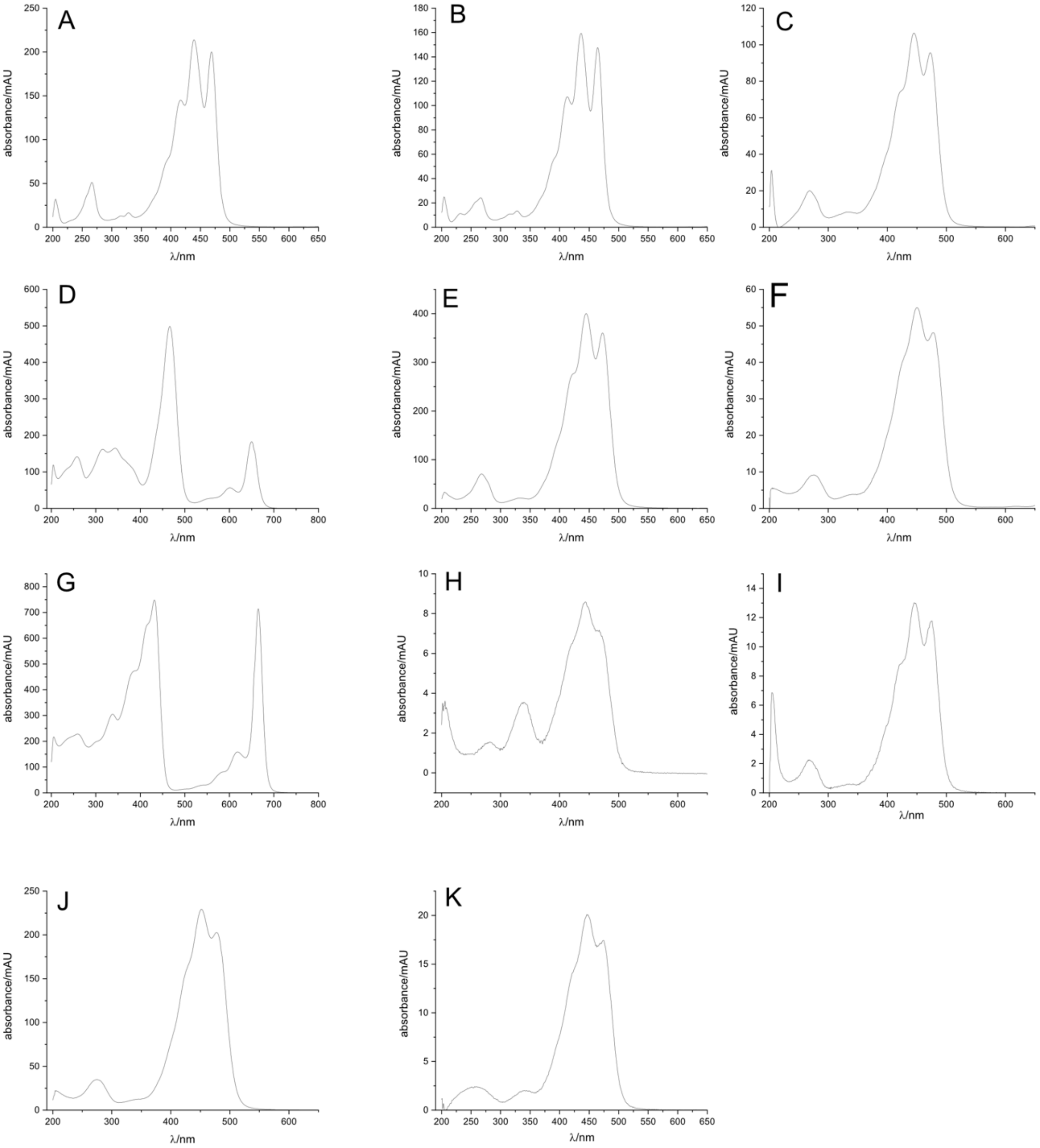


**Supplemental Figure 3:** Absorption spectra extracted from *Mesotaenium endlicherianum* control replicate 1. Violaxanthin (A), 9-*cis*-neoxanthin (B), antheraxanthin (C), chlorophyll b (D), lutein (E), zeaxanthin (F), chlorophyll a (G), 15-*cis*-β-carotene (H), α-carotene (I), β-carotene (J), 9-*cis*-β-carotene (K).

**Supplemental Table 2: Calibration curves of carotenoid and chlorophyll standards**

| Pigment (retention time/min) | Effective amounts of calibration applied to column/µg | Linear regression (y = µg; x = mAU) | Detected absorption maxima (reported maxima [Gupta et al 2015])/ nm |
|---|---|---|---|
| violaxanthin (4.38) | 0.0257, 0.257, 1.286, 2.571 | $y = 6.021 \cdot 10^{-4}x$ | 266, 416.5, 439, 469 (416, 439, 469) |
| 9-*cis*-neoxanthin (4.58) | see violaxanthin | see violaxanthin and text below | 266, 328, 413, 436, 465 (415.0, 437.0, 465.0) |
| antheraxanthin (5.60) | see zeaxanthin | see zeaxanthin and text below | 268, (423), 445.5, 473.5 (424, 446, 474) |
| chlorophyll b (Milenković et al 2012) (5.82) | 0.0271, 0.271, 1.357, 2.714 | $y = 5.897 \cdot 10^{-4}x$ | 344, 466, 601, 650 (461, 598, 648) |
| lutein (6.46) | 0.0286, 0.286, 1.429, 2.857 | $y = 5.949 \cdot 10^{-4}x$ | 268, (423), 445, 473 (424, 445, 474) |
| zeaxanthin (7.32) | 0.0257, 0.257, 1.286, 2.571 | $y = 2.646 \cdot 10^{-4}x$ | 276, (428), 450.5, 478 ((428.0), 451, 478) |
| chlorophyll a (Milenković et al 2012) (7.78) | 0.0271, 0.271, 1.357, 2.714 | $y = 2.577 \cdot 10^{-4}x$ | 338, (383), (413), 432, (533), (581), 619, 665 (411, 431, 532, 581, 617, 662.5) |
| 15-*cis*-β-carotene (11.55) | see β-carotene | see β-carotene | 339, (424), 444, 467 (338.0, (420.0), 444.0, 468.0) |
| α-carotene (11.81) | 0.0271, 0.271, 1.357, 2.714 | $y = 3.719 \cdot 10^{-4}x$ | 266, (420), 445.5, 475 (424, 446, 475) |
| β-carotene (12.62) | 0.0266, 0.266, 1.329, 2.657 | $y = 0.00141x$ | 275, (426), 452, 478 ((425.0), 452, 479) |
| 9-*cis*-β-carotene (13.26) | see β-carotene | see β-carotene | 256, 340, (421), 446, 473.5 ((420.0), 447.0, 473.0) |
| lycopene (19.42) | 0.00643, 0.0643, 0.321, 0.643 | $8.739 \cdot 10^{-5}x$ | 295, 445, 472, 503 (446.0, 472.0, 503.0) |

**Supplemental Table 2: Quotients of extinction coefficients used for calibration**

| Pigment | Extinction coefficient based on Thrane et al 2015 / L $g^{-1}$ $cm^{-1}$ | Quotient of extinction coefficients ($E_2/E_1$) |
|---|---|---|
| 9-*cis*-neoxanthin ($E_2$) | 233 | 0.9173 |
| violaxanthin ($E_1$) | 254 | |
| antheraxanthin ($E_2$) | 235 | 0.9592 |
| zeaxanthin ($E_1$) | 245 | |

Retention times of analytical standards and plant samples differed slightly due to effects of the plant matrix (and so also between organisms). Retention times based on *Mesotaenium endlicherianum* control replicate 1 except lycopene (based on standard), which did not accumulate under our conditions. The pistons appeared to be subject to relatively high loads, so that the retention times changed over time and the pistons had to be replaced more frequently than usual. Quality of analytical standards of vendors was strongly varying although treated with care (stored at –80 °C, weighted for calibration in the dark, measurements started on the same day as stock preparation etc.). Some were not mono isomeric and in the case of 9-*cis*-neoxanthin standard quality was so low that violaxanthin was used for calibration respecting differences in reported extinction coefficients (see table above). Antheraxanthin was quantified based on zeaxanthin calibration (since no commercial standard was at hand at this time) respecting the differences in reported extinction coefficients (see table above).

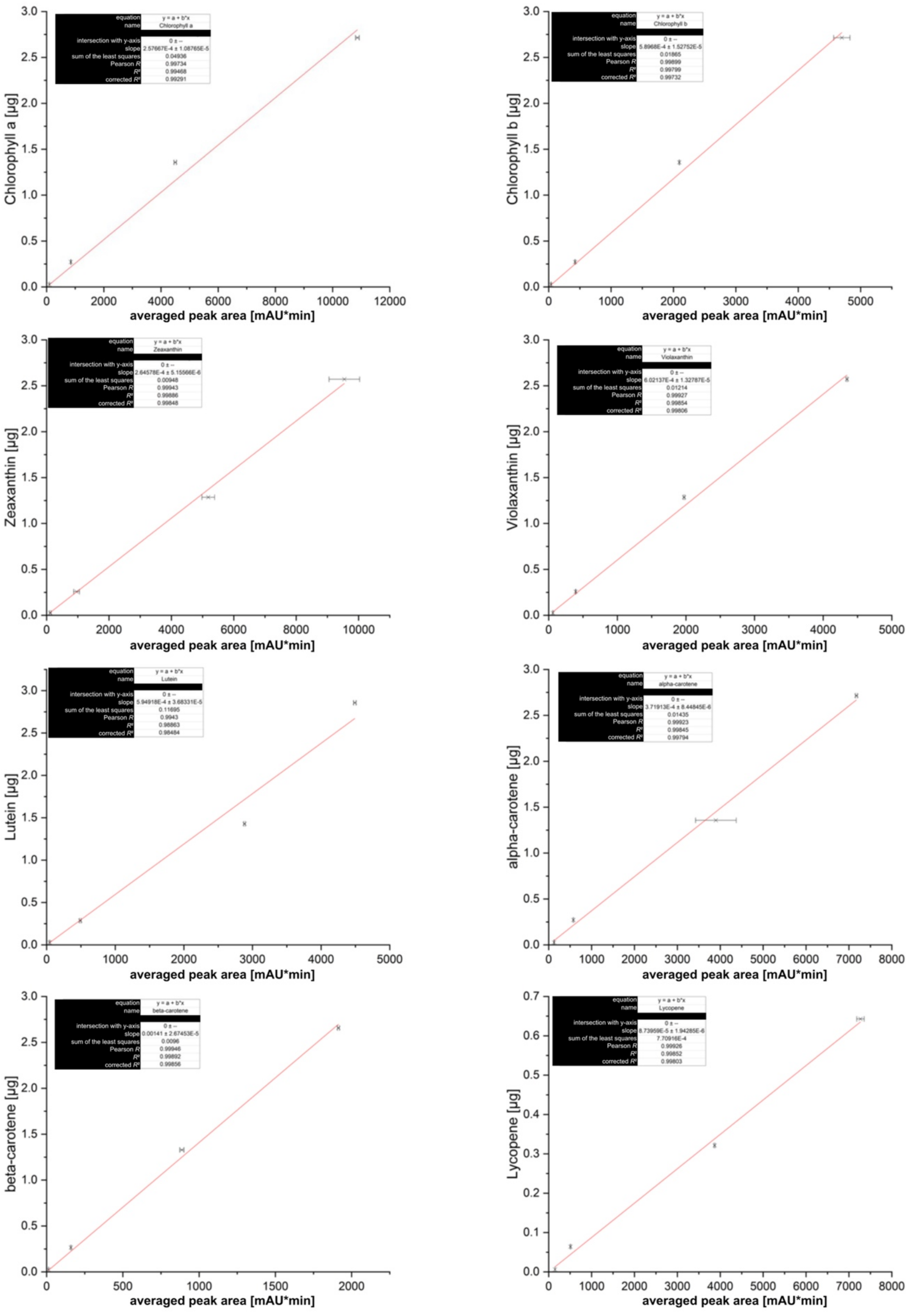


**Supplemental Figure 4:** Linear regressions of carotenoid and chlorophyll calibration data based on authentical analytical standard dilutions. X-axis: Averaged peak area of triplicate measurements in mAU*min; Y-axis: Respective metabolite and total amount of injected standard in µg.

### HS-SPME-GC-MS

#### Recovery rate

HS-SPME methods do come with matrix effects during metabolite adsorption (and minor effects during desorption), either influenced by one or several metabolites or the complex matrices created by the biomass of the organisms. To normalize results the recovery rate of coinjected β-ionone-$D_3$ (4 µL) was applied. To determine this rate the measurements of the calibration (with β-ionone-$D_3$ coinjection) of 6-methyl-5-hepten-2-one (in total 9 replicates), the apocarotenoid standard with the lowest matrix effect on the β-ionone-$D_3$ signal (no significant effects) were used. The average signal of β-ionone-$D_3$ was used as reference ($R_{\beta\text{-ionone-D3}}$) for normalization and every signal detected was multiplied by the resulting recovery rate comparing the actual β-ionone-$D_3$ signal of each measurement with $R_{\beta\text{-ionone-D3}}$

$$\text{recovery rate} = \frac{R_{\beta-\text{ionone}-\text{D3}}}{\text{signal } \beta-\text{ionone}-\text{D3}}$$

**Supplemental Table 3: Calibration curves of apocarotenoid standards**

| Apocarotenoid (m/z: **quantifying ion**, second qualifier, retention time) | Effective amounts of calibration applied to column/ng | Linear regression (y = ng; x = AU) |
|---|---|---|
| 6-methyl-5-hepten-2-one (**108**, 126, 6.12) | 24, 120, 240 | $y = 7.107 \cdot 10^{-6}x$ |
| β-cyclocitral (**137**, 152, 12.32) | 0.522, 1.044, 5.22, 26.1, 52.2 | $y = 1.562 \cdot 10^{-6}x$ |
| β-ionone (**177**, 192, 19.20) | 0.101, 0.202, 1.012 | $y = 1.241 \cdot 10^{-8}x$ |
| dihydroactinidiolide (**111**, 180, 20.19) | 0.152, 0.304, 1.522, 7.612, 15.224 | $y = 4.087 \cdot 10^{-7}x$ |

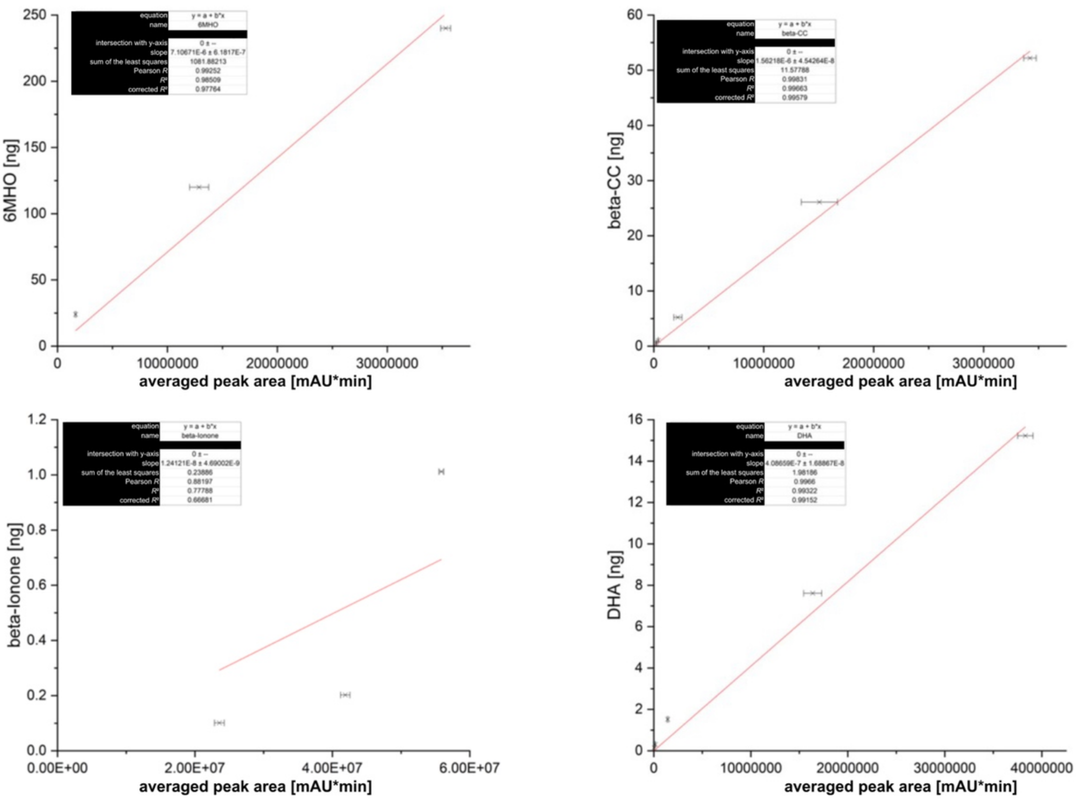


**Supplemental Figure 5:** Linear regressions of apocarotenoid calibration data based on authentical analytical standard dilutions. X-axis: Averaged peak area of triplicate measurements in mAU*min; Y-axis: Respective metabolite and total amount of injected standard in ng.

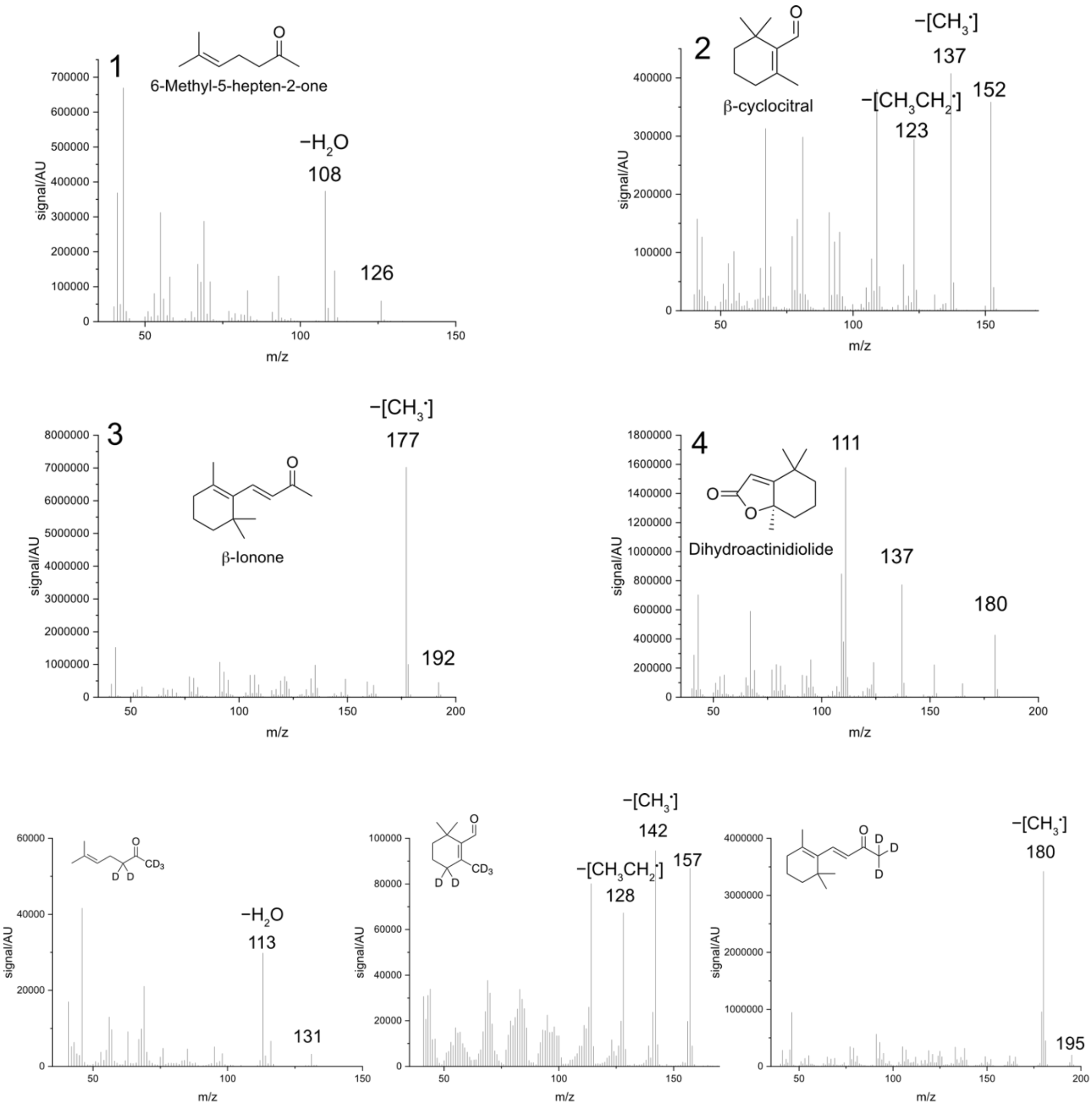


**Supplemental Figure 6:** Fragmentation patterns of authentical commercial standards and deuterated standards. Top: authentical commercial standards. Spectrum 1 shows the fragmentation pattern of 6MHO, with the molecular mass of 126 and the fragment obtained after loss of water (108). Spectrum 2 shows the fragmentation pattern of beta-cyclocitral with a molecular mass of 152, a fragment after loss of a methyl group (137) and a fragment that has lost an ethyl / two methyl groups (123). Spectrum 3 shows the fragmentation pattern of beta-ionone with a molecular mass of 192, and a fragment that has lost a methyl group (177). Spectrum 4 shows the fragmentation pattern of DHA with a molecular mass of 180, and two characteristic fragments of 137 and 111. Qualifying ions were picked based on literature, including Ramel et al. (2012) and Rivers et al. (2019). Bottom: deuterated standards.

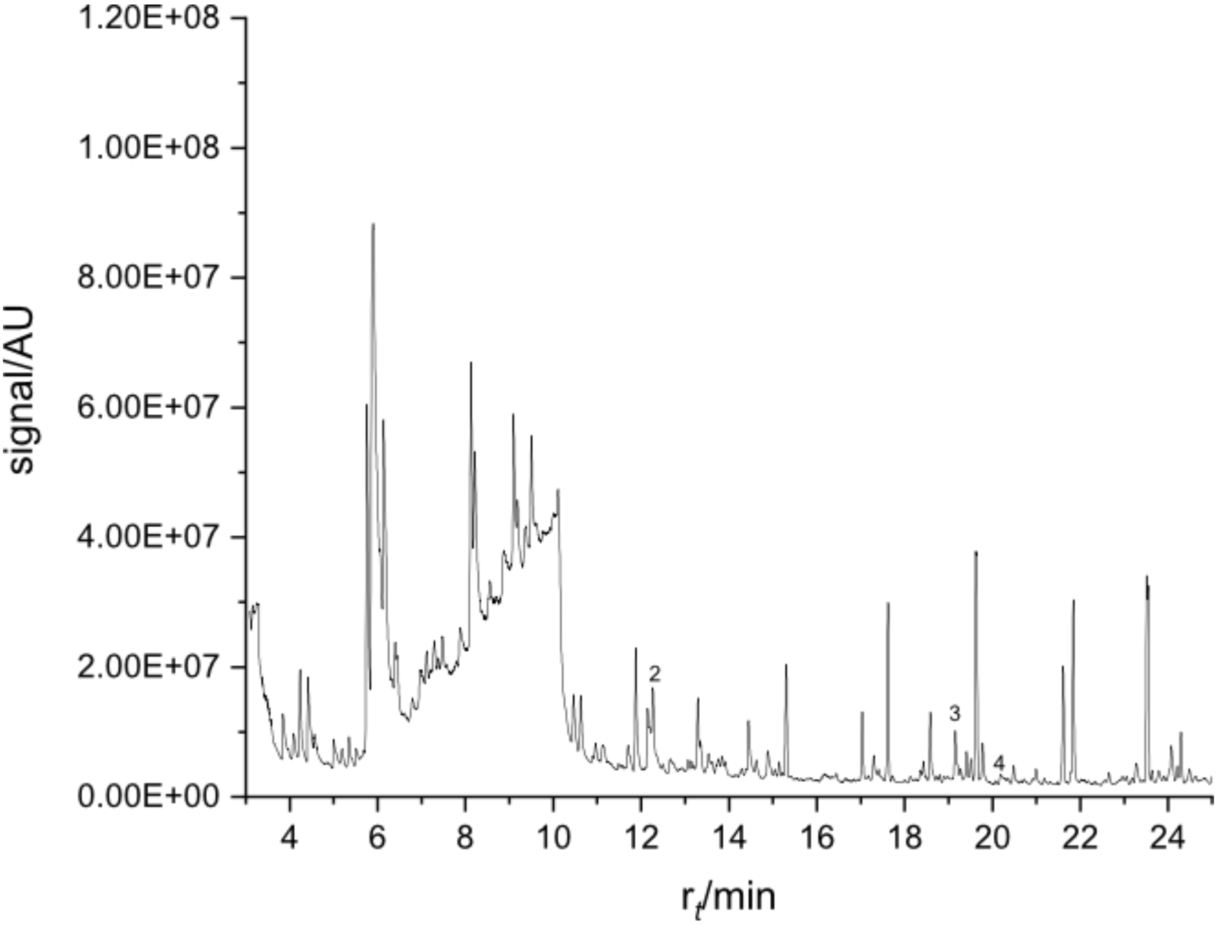


**Supplemental Figure 7**: Total ion chromatogram (TIC) of *Physcomitrium patens*. Homogenized flash frozen and lyophilized plant material was heated and volatiles were absorbed at the SPME fiber prior to injection and desorption. Detection was carried out using the scan mode of the mass spectrometer over a measuring time of 34.4 minutes; all relevant signals salient to apocarotenoids eluted before 24 minutes.

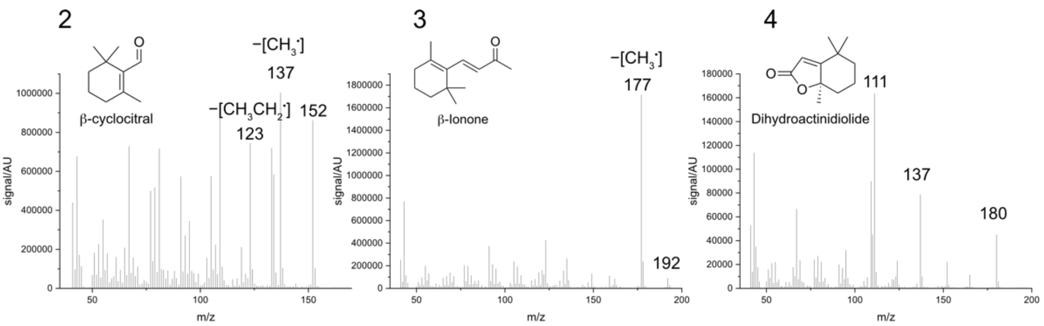


**Supplemental Figure 8**: Fragmentation pattern of apocarotenoids in *Physcomitrium patens*. Detection was based on informative fragmentation patterns as described for the commercial standards (see Supplemental Fig. 6).

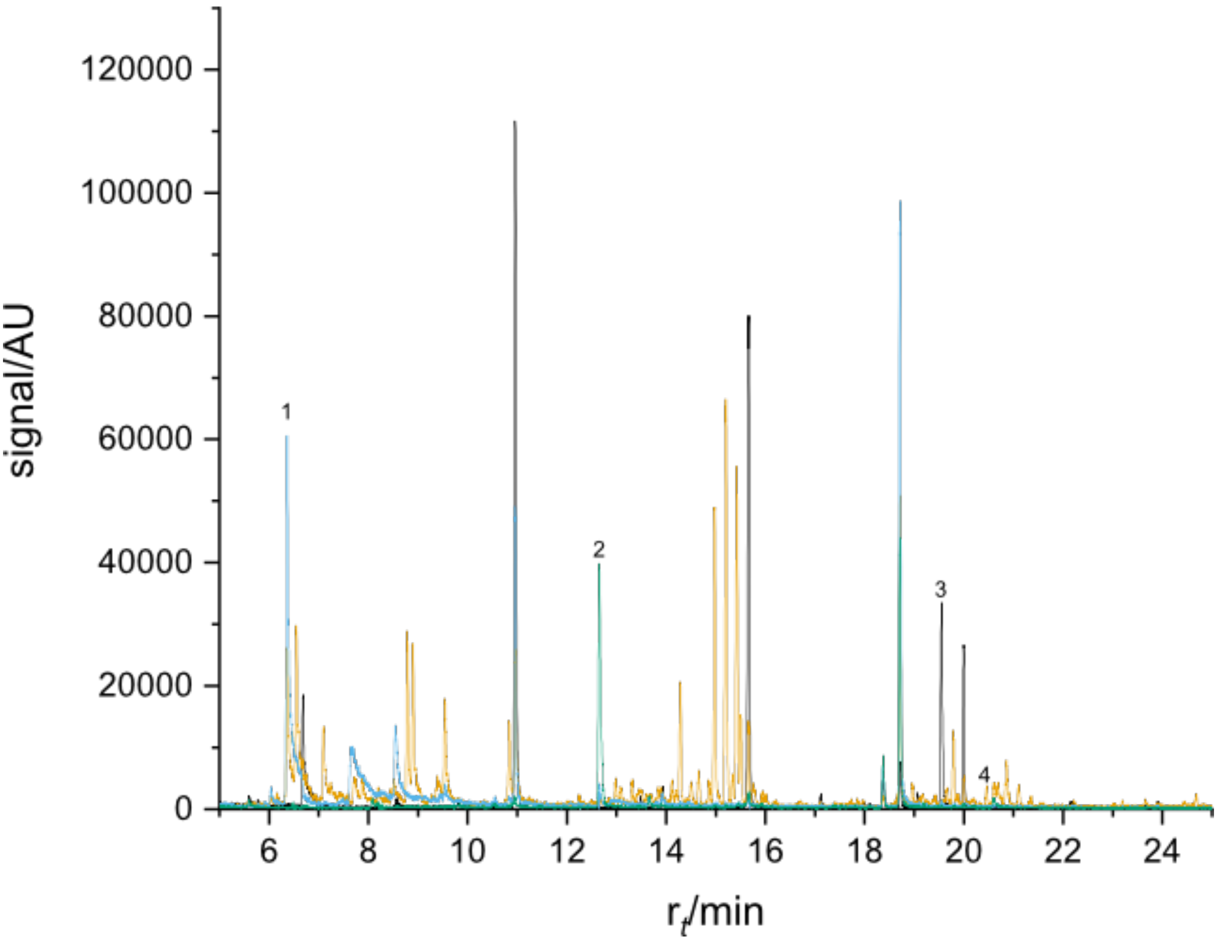


**Supplemental Figure 9**: Selected and merged ion chromatograms (XIC) of *Mesotaenium endlicherianum*. Measurements were performed in scan mode yielding TICs and later, during data analysis, selected ion chromatogram were merged to yield XICs. Homogenized flash frozen and lyophilized plant material was heated and volatiles were absorbed at the SPME fiber prior to injection and desorption. Detection was carried out using the scan mode of the mass spectrometer over a measuring time of 34.4 minutes; all relevant signals salient to apocarotenoids eluted before 24 minutes.

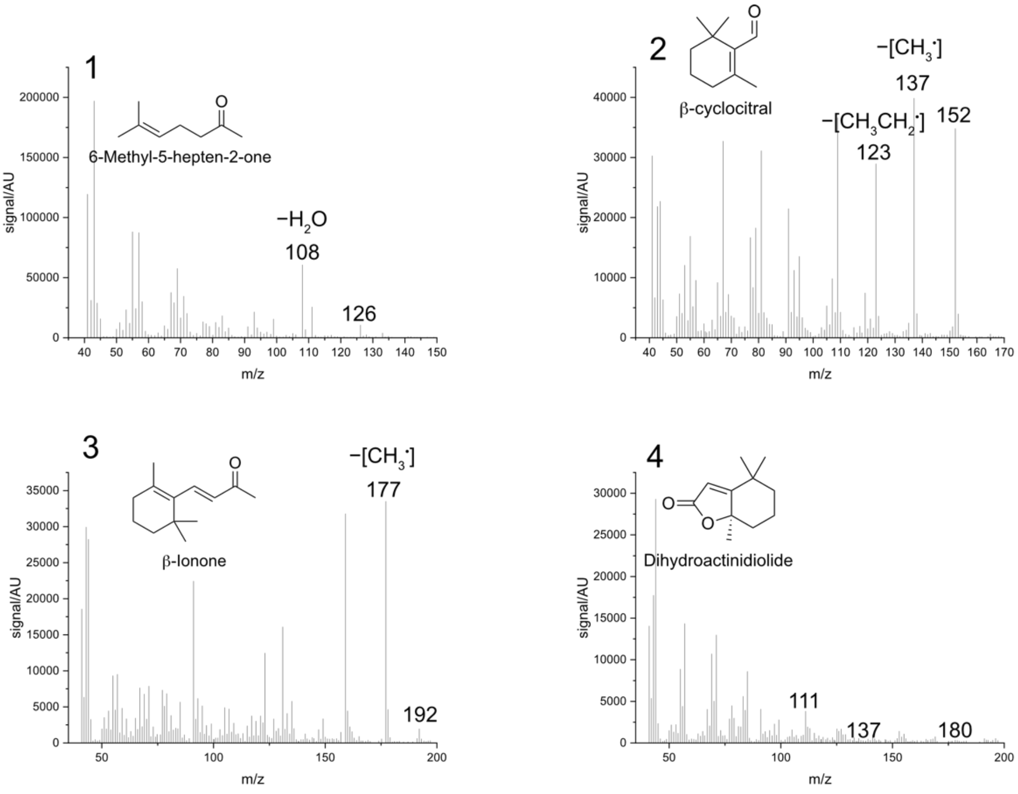


**Supplemental Figure 10**: Fragmentation pattern of apocarotenoids in *Mesotaenium endlicherianum*. Detection was based on informative fragmentation patterns as described for the commercial standards (see Supplemental Fig. 6).

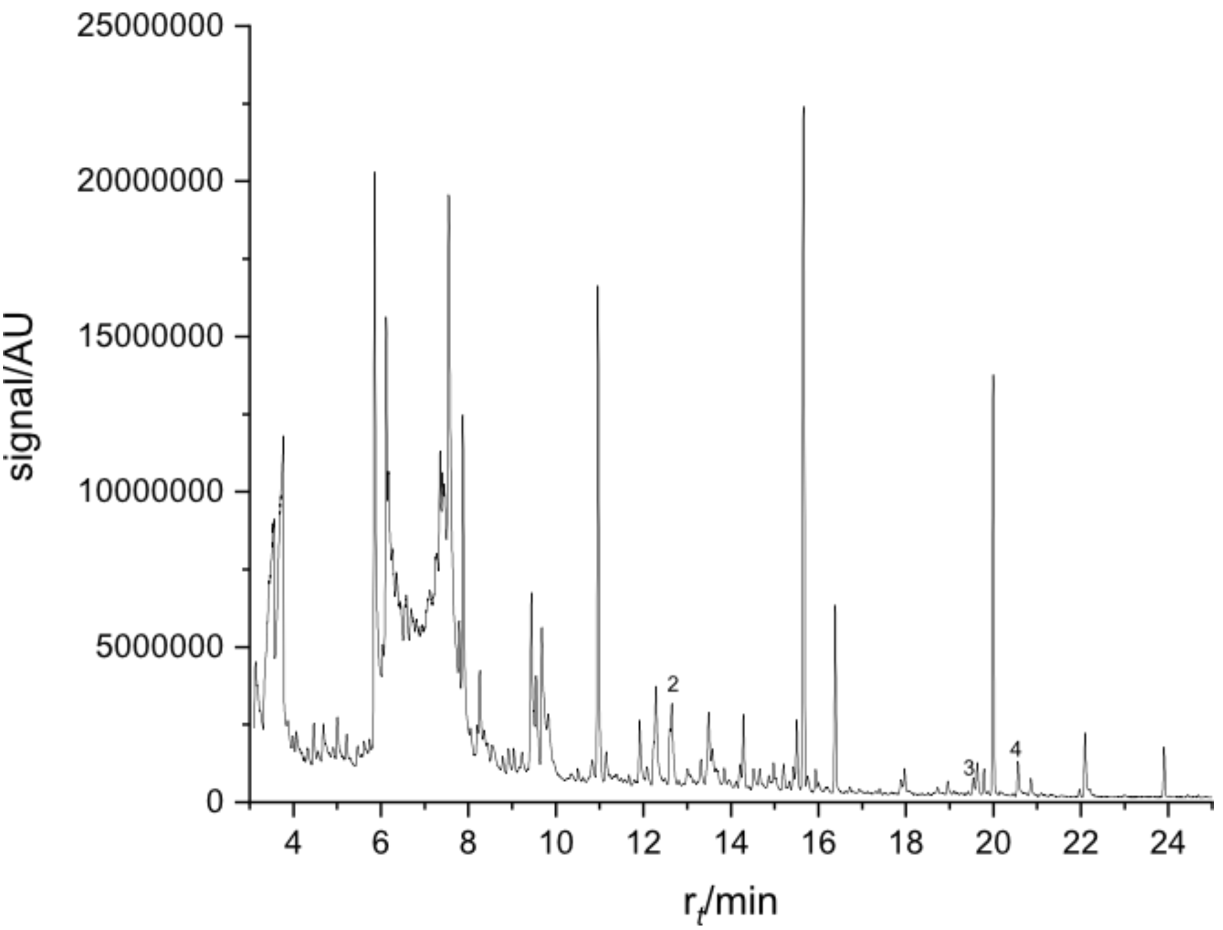


**Supplemental Figure 11**: Total ion chromatograms (TIC) of *Zygnema circumcarinatum*. Homogenized flash frozen and lyophilized plant material was heated and volatiles were absorbed at the SPME fiber prior to injection and desorption. Detection was carried out using the scan mode of the mass spectrometer over a measuring time of 34.4 minutes; all relevant signals salient to apocarotenoids eluted before 24 minutes.

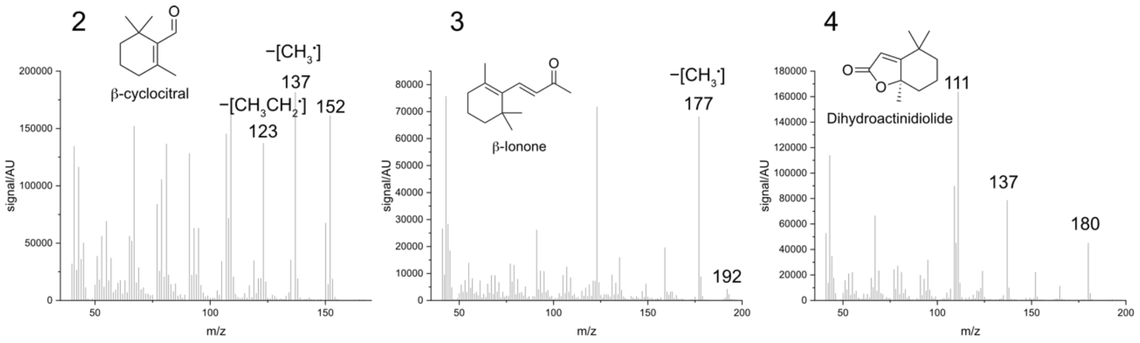


**Supplemental Figure 12**: Fragmentation pattern of apocarotenoids in *Zygnema circumcarinatum*. Detection was based on informative fragmentation patterns as described for the commercial standards (see Supplemental Fig. 6).
